# CuBi-MeAn Customized Pipeline for Metagenomic Data Analysis

**DOI:** 10.1101/2021.09.10.458355

**Authors:** Saeed Keshani-Langroodi, Christopher M. Sales

## Abstract

Whole genome shotgun sequencing is a powerful to study microbial community is a given environment. Metagenomic binning offers a genome centric approach to study microbiomes. There are several tools available to process metagenomic data from raw reads to the interpretation there is still lack of standard approach that can be used to process the metagenomic data step by step. In this study CuBi-MeAn (Customizable Binning and Metagenomic Analysis) create a customizable and flexible processing pipeline, to process the metagenomic data and generate results for further interpretation.

This study aims to perform metagenomic binning to enhance taxonomical classification, functional potentials, and interactions among microbial populations in environmental systems. This customized pipeline which is comprised of a series of genomic/metagenomic tools designed to recover better quality results and reliable interpretation of the system dynamics for the given systems. For this reason, a metagenomic data processing pipeline is developed to evaluate metagenomic data from three environmental engineering projects.

The use of our pipeline was demonstrated and compared on three different datasets that were of different sizes, from different sequencing platforms, and generated from three different environmental sources. By designing and developing a flexible and customized pipeline, this study has showed how to process large metagenomic data sets with limited resources. This result not only would help to uncover new information from environmental samples, but also, could be applicable to any other metagenomic studies across various disciplines.

## 2. Introduction

Advances in molecular biology techniques such as next generation sequencing (NGS), PCR, molecular cloning, DNA Microarrays and protein mass spectrometry have improved our knowledge of microbiomes (Jovel, Patterson et al. 2016, Mendes, Braga et al. 2017, Quince, Walker et al. 2017, Heyer, Schallert et al. 2019, Sun, Liao et al. 2020). Among them, NGS is becoming one the most popular techniques used to generate targeted amplicon sequencing, shotgun metagenomics, and meta-transcriptomics data to study microbial communities. NGS technologies depend on sequencing technologies capable of generating millions (and sometimes billions) of small fragments of short (e.g. Illumina and SOLiD) or long (e.g. PacBio and Oxford Nanopore) DNA sequences (Mardis 2008, Quail, Kozarewa et al. 2008, Rhoads and Au 2015, Jain, Olsen et al. 2016). These NGS technologies can be used to produce sequencing libraries which contain substantial information about the entire microbial population living in an environment. Depending on the research question, scientists may choose between two type of popular approaches for generating libraries from DNA isolated from samples: amplicon sequencing and metagenomic sequencing. Amplicon sequencing analysis is aimed at exploring microbial community composition based on targeted sequencing of PCR amplicons of certain conserved region (16S rRNA, ITS or 18S) of the genomes that could be used as unique marker for phylogenic classification of the organisms present in a sample. Since this method relies on existing databases for classifications, they are inherently biased towards known organisms (Walsh, Crispie et al. 2018). In addition, taxonomical classification based on only marker genes fails to address the functional variation within closely related genomes (Hiergeist, Gläsner et al. 2015, Tremblay, Singh et al. 2015, Gohl, Vangay et al. 2016). Isolated DNA from a mixed microbial community can also be used for metagenomic shotgun sequencing, which has become increasingly utilized to characterize both taxonomy and function of microbial communities (Woloszynek, Zhao et al. 2018). Through metagenomic shotgun sequencing, it is possible to generate sequence libraries containing the genetic information from hundreds or even millions of cells in sample, in order to understand their taxonomical classification, potential functional capabilities, and physiological traits. Metatranscriptomic is another approach reliant on NGS technologies commonly used how genes are regulated in response toward environmental factors and stimuli (Ranjard, Poly et al. 2000). The rest of this chapter only focuses on developing a pipeline for processing the metagenomic data and producing outputs that could be used in downstream analysis and interpretations such as taxonomical classification, interactions and potential functions of the microbial communities exist in a microbiome. For more information on targeted amplicon sequencing and meta-transcriptomics, please see Rausch et al. (2019) article and Hodkinson and Grice (2015) and Bashiardes et al. (2016) reviews.

Recovering information from several fragments of DNA sequences generated by NGS facilities (also known as “reads”) is not an easy task (Tyson, Chapman et al. 2004). Currently, there are two approaches for processing metagenomic libraries: gene-centric and genome-centric approaches. Gene-centric approaches (Venter, Remington et al. 2004, Tringe, Von Mering et al. 2005) involve the recovery and investigation of the entire microbiome as a “supra-organism” regardless of their individual function (Juengst and Huss 2009, Juengst 2009). In gene-centric approaches, individual genes are regarded as selfish units and are the central keys in carrying out the functions while genomes are nothing more than the vessel for the genes (Dawkins 2016). Here, the genes are the fundamental framework of molecular biology for decoding the blueprint of the life and evolution (Venter, Adams et al. 2001, Tishkoff and Verrelli 2003, Schloss and Handelsman 2004, Guénet 2005). Gene-centric approaches rely heavily on existing databases and often overlook novel genes (Jaenicke, Ander et al. 2011, Wong, Zhang et al. 2013). In gene centric approach certain functions are attributed to a gene or a gene cluster. These genes are going to be used as a reference for annotation of the unknown genes. Therefore, any variation of these genes may increase the errors of annotations. In addition, another problem is confusion of homologous genes that have very similar genes may have different functions. For example, ammonia monooxygenases is very similar to methane monooxygenases. This could cause perplexity to annotation and interpretation of annotation. In addition, especially for short read sequencing libraries, this approach fails to address questions related to the function of individual genes without considering that metabolic and functional traits could be dependent on multiple genes and how they are regulated. For example, in metagenomic investigation approaches, a certain pathway is complete in gene-centric approaches, however, in this approach it remains unspecified if all of the genes belong to one organism or belong different organisms. For proper function of the some pathways intermediates/metabolites may need to be transported out and into the cell (Strambio-De-Castillia, Niepel et al. 2010, Villegas and Zaphiropoulos 2015). This would be also true about the co-expression or co-regulation of the genes. Cells respond to environmental changes by reprogramming expression of specific genes throughout the genome. The transcription rate of a particular gene is determined by the interaction of diverse regulatory proteins—transcriptional activators and repressors—with specific DNA sequences in the gene’s promoter. How a collection of regulatory proteins accomplishes the task of regulating a set of genes can be described as a regulatory network (Wyrick and Young 2002). These networks might be present in a dataset, but the arrangement or transporters control the proper function of these genes. Therefore, in this approach metabolic interactions of genes is difficult to prove (Heng 2009, Vanwonterghem, Jensen et al. 2016).

These shortcomings in gene centric approaches in metagenomic studies led to the development of the genome-centric concept which has revealed the functional properties of individual genomes, leading to a more detailed comprehension of the microbial interactions occurring in the microbiome (Kougias, Campanaro et al. 2018). While, gene-centric approaches focus on to the function of individual genes and correlating it to the biochemical activities of the system, genome-centric approaches decipher the complexity of the genome by considering the genes functions and interplay within a genome. Genome-centric approaches create an additional dimension to functional analysis of metagenomic data by correlating the interaction of the products of the different genes existing in a genome and environmental factors (Raghoebarsing, Pol et al. 2006, Wrighton, Castelle et al. 2014, Brown, Hug et al. 2015, Castelle, Wrighton et al. 2015).

The genome-centric concept is based on the premise that a microbial community is composed of taxonomically and functionally related bacterial populations that can interact. Each bacterial population is comprised of a ‘core-genome’, consisting of genes that are always present and carry out major functions and a ‘pan-genome’ which contains genes that are variably present (Tettelin, Masignani et al. 2005). The pan-genome is a holistic snapshot of the collective genomes from closely related organisms and thus includes specific and specialized functions and adaptations of divergent taxonomical units belonging to the diversity among species or strains that compose the pan-genome. This provides valuable genetic data for understanding the evolutionary processes which affect the structure and dynamics of related bacterial populations in relation to the environmental factors (Holmes, Gillings et al. 2003, Whitaker and Banfield 2006). Furthermore, integrating functional and taxonomical results by using genome-centric methods and coupling them to the existing databases enables us to have a deeper and more comprehensive insight into dynamics of a biological system.

While gene-centric analysis is heavily dependent on assembling fragmented DNA reads, genome centric analysis of metagenomic data depends on the clustering of reads into bins. “Binning” is a method for clustering reads based on certain characteristics which is used as an alternative to full metagenome assembly which is the basically assembly of the reads to a supra-organism for downstream analysis (Teeling, Waldmann et al. 2004, Woyke, Teeling et al. 2006, Albertsen, Hugenholtz et al. 2013, Cotillard, Kennedy et al. 2013, Le Chatelier, Nielsen et al. 2013, Alneberg, Bjarnason et al. 2014). Development of new algorithms improved the metagenomic tools used in processing metagenomic data including binning tools (Anantharaman, Brown et al. 2016, Parks, Rinke et al. 2017, Almeida, Mitchell et al. 2019, Pasolli, Asnicar et al. 2019). There are different metagenomic binning tools available for processing metagenomic data. These metagenomic binning tools use K-mer frequency, codon content, and read coverages read coverages across multiple data sets for clustering the short read into the bins (Alneberg, Bjarnason et al. 2014, Kang, Froula et al. 2015, Wu, Simmons et al. 2015, Graham, Heidelberg et al. 2017, Lu, Chen et al. 2017). However, due to different algorithms used in these tools, generated bins could be different for the same data bases. Therefore, refining tools is developed intended to improve the quality of the quality of the bins (Sieber, Probst et al. 2018). These improved quality bins can be used in taxonomical classifications to approximate the position of bins into the phylogenetic tree and the potential metabolic pathways that these bins could carry out.

There are many methods to go from raw reads to bins but there are no methods that make use of these bins to making use of bins to gain information about the function of different populations identified in a microbial community and how they may, interact. Therefore, the purpose of the pipeline described here is to go from processing of the metagenomic data from starting point to take the sequencing data and generate data that are ready for downstream analysis and interpretations. Although, there are many powerful tools available to process metagenomic data from raw reads to the interpretation there is still no standard approach that can be used to process of a standard approach that user could be used to process the metagenomic data step by step. Existing cloud services have some limitation such as the size of the data bases high dependency on internet and lack of flexibility of the options. Uritskiy et al. (2018) developed a pipeline to process metagenomic data in this pipeline which is called MetaWrap. MetaWrap is an automated pipeline which comprised of several metagenomic tools that process raw reads from metagenomic samples, cluster them into the metagenomic bins then generate outputs for final interpretation such as taxonomical classification and functional annotations. This pipeline includes several tools that all required to be installed and run to generate the results however, user would be able to customize individual tools, but the overall processing steps are almost fixed and need to be run.

Another metagenomic data processing and interpretation pipeline created by Clarke et al. (2019) named Sunbeam. This pipeline includes series of metagenomic tools for quality control, decontamination assembly, taxonomical classification and functional annotations. Unlike MetaWrap, Sunbeam uses pre-processed metagenomic reads for taxonomical classification rather than clustering the bins into “genome”. This tool used mapped reads for functional annotation. The advantage of this pipeline is due to the parallel configuration of tools which make the steps independent and the pipeline highly flexible and customizable. In this pipeline taxonomical classification and annotation are directly from the pre-processed reads and independent of each other. Therefore, the user will not be able to investigate assigned functions and taxonomy at the same time to correlate interaction of the genes within a genome which is the basic of genome centric approach.

The aim of this work is to create a more customizable and flexible processing pipeline to process the metagenomic data and generate results for further interpretation. This pipeline which is called CuBi-MeAn (Customizable Binning and Metagenomic Analysis) generates taxonomical classification and functional annotations that could be used for genome-centric as well as gene-centric investigation of the given microbiome. CuBi-MeAn is comprised of a series of metagenomic tools that could be customized by user. The flexibility of this pipeline allows the users to add new tools to each step (ex. different assembly or binning tools can be added). Since, the tools in CuBi-MeAn are independently installed the user would be able to install and use them on separate system such as shared clouds or local systems. This flexibility would be an advantageous when users handling large size metagenomic datasets which need to deal with some system limitations (RAM, storage, etc.). In the following section we reviewed the details, different steps and the tools used in CuBi-MeAn; Then, we discussed about the performance of this pipeline on processing three different metagenomic datasets.

## 3. Methodology

CuBi-MeAn is comprised of a series of metagenomic tools that are able to use metagenomic raw reads as input and generate bins for functional annotation and taxonomical classifications. The overall workflow is summarized in Figure 2.1. The modules used in this pipeline may require different dependencies that could be installed and run separately. The Anaconda installation package and module instruction is also available for most of the modules. Detailed instruction and functions are covered in the following sections.

**Figure 2. 1:**
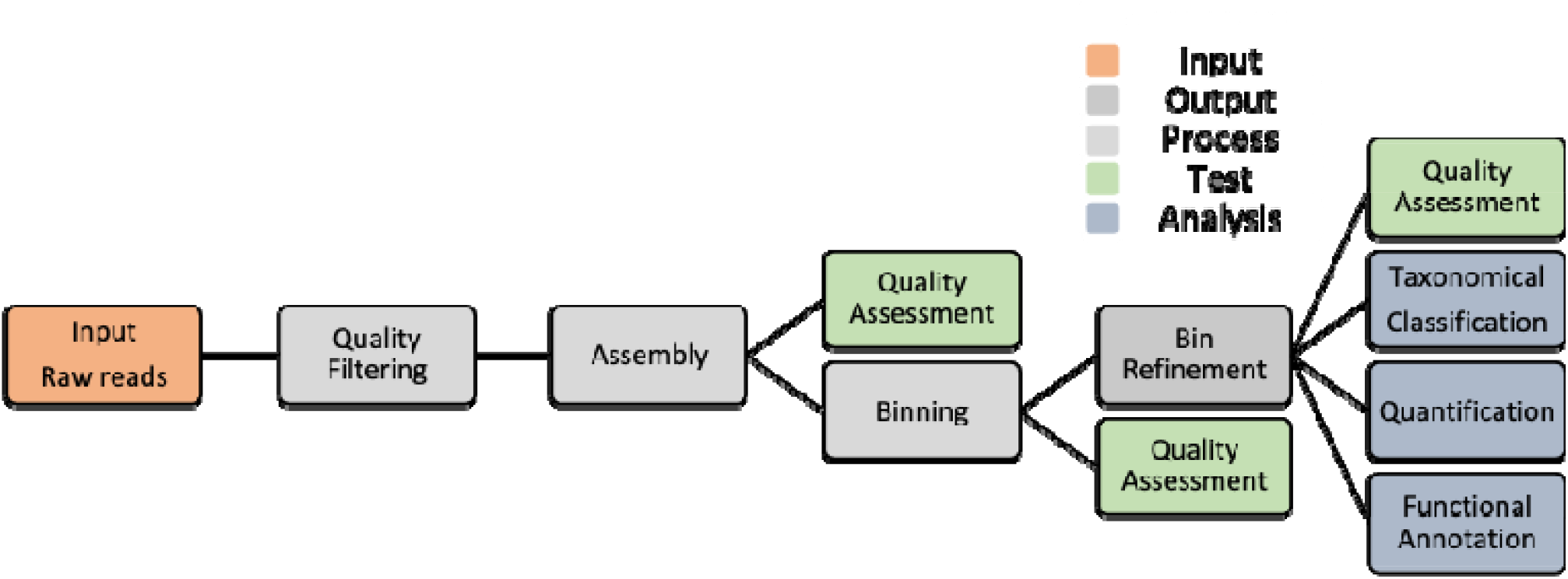
CuBi-MeAn overall workflow.

### 1.1. Data preparation

#### Quality Filtering

To improve the quality of the raw data, Sickle tool is used to remove low quality end of the reads (Joshi and Fass 2011). Users can customize the options of Sickle tool based on sequencing technology and input and output (https://github.com/najoshi/sickle).

#### Input preparation

For assembly tools (merge paired-end reads, convert fastq to fasta, etc). Several tools are available for this purpose such as FASTXToolkit (http://hannonlab.cshl.edu/fastx_toolkit/commandline.html) or IDBA package tools (https://github.com/loneknightpy/idba).

##### 1.1.1. Data processing

###### Assembly

Assembly of the entire metagenomic libraries is the first step for the metagenomic binning. De-novo metagenomic assembly tools which does not require reference genomes are used for assembly of the raw reads into the contigs.

Here, IDBA de-novo assembly tool was used to assemble the whole metagenomic data (Peng, Leung et al. 2010). In this step, the whole metagenomic libraries are assembled using appropriate metagenomic tools to create a “Supra-genome”. This “supra-genome” which represent the entire metagenomes use as “reference” for the next binning clustering step. Initially, MetaVelvet (Namiki, Hachiya et al. 2012), IDBA (Peng, Leung et al. 2010) and Celera Assembler (Myers, Sutton et al. 2000, Venter, Adams et al. 2001) were tested for their performance which has been evaluated by certain parameters such as contigs size, N50, etc. To evaluate the quality of assembled metagenomes “summarizeAssembly.py” tool from PBSuite package was used (https://github.com/dbrowneup/PBSuite).

Then, selected assembly tools were compared for their computational resources’ requirements. Among them IDBA outperformed the others used for metagenomic assembly in this pipeline. The user may customize IDBA parameters to optimize the assembly (https://github.com/loneknightpy/idba).

###### Binning

In this study, the metagenomic binning approach was utilized to investigate the subject metagenomic data sets. The two different main approaches for binning of metagenomic data are supervised and unsupervised binning. In supervised binning, in reference genomes are used for clustering of the metagenomic reads. Supervised binning is suitable when there are specific targeted species in the data set. In this method the reference genomes are aligned to the query and binning is based on the GC content and k-mers frequency (Mohammed, Ghosh et al. 2011, Mande, Mohammed et al. 2012). However, the accuracy of the supervised methods is questionable, especially for environmental samples that have higher diversity (Sedlar, Kupkova et al. 2017). In addition, supervised method could be biased towards the reference genomes and leave out new species (Cole, Brosch et al. 1998).

Another binning approach is called unsupervised binning in which the reference genome is not required for read clustering. Instead, unsupervised binning relies on sequence composition, abundance of genome fragments or a hybrid method. Nucleotide composition methods are based upon a theory that oligonucleotide, dinucleotide or G+C content showed species-specific pattern within the DNA of the same genomes (Sandberg, Winberg et al. 2001, Pride, Meinersmann et al. 2003, Wu and Ye 2011). This notion became the main pillar for design of the algorithm of tools such as TETRA (Teeling, Waldmann et al. 2004), MetaCluster (Woyke, Teeling et al. 2006), and MetaCluster, (Yang, Peng et al. 2010). However, there are still some disputes on accuracy of the sequence composition method due to sequence variation within a single genome, which makes it challenging to accurately classify very short reads (Yang, Peng et al. 2010, Wu and Ye 2011). It has been suggested that the abundance of certain genes (aka marker genes) are constant across the same genomes (Wang, Leung et al. 2012, Albertsen, Hugenholtz et al. 2013, Nielsen, Almeida et al. 2014). In the other word, for a given sample the abundance of a gene in a specific genome will be the same as other genomes of the same species. Therefore, coverage-based binning was introduced as an alternative for composition-based binning in unsupervised binning (Albertsen, Hugenholtz et al. 2013, Cotillard, Kennedy et al. 2013, Le Chatelier, Nielsen et al. 2013, Alneberg, Bjarnason et al. 2014). These two principals were later integrated to create hybrid binning tools such as BinSanity (Graham, Heidelberg et al. 2017), MaxBin2 (Wu, Simmons et al. 2015), MetaBAT (Kang, Froula et al. 2015), COCACOLA(Lu, Chen et al. 2017) and CONCOCT (Alneberg, Bjarnason et al. 2014) that outperformed any of those individual tools. Therefore, unsupervised hybrid binning tools are ideal option for metagenomic analysis of samples with higher diversity such as environmental samples.

In this study, metagenomic assembled contigs from the assembly step were used as the reference for binning tools. CuBi-MeAn utilize the following five hybrid binning tools: BinSanity (Graham, Heidelberg et al. 2017), MaxBin2 (Wu, Simmons et al. 2015), MetaBAT (Kang, Froula et al. 2015), COCACOLA (Lu, Chen et al. 2017) and CONCOCT (Alneberg, Bjarnason et al. 2014). As these tools are able to be run in parallel within this pipeline in this pipeline, the user capable of add new binning tools or opt out any aforementioned binning tools. More information regarding these binning tools can be find in their refence and webpages.

###### Bins refinement

The results of binning tools can be used for downstream analysis. However, since many of these binning tools use different parameters and approaches (i.e. different algorithms) for processing metagenomic data; This results low quality and incomplete bins and better performance for different data sets using the same binning tool. Therefore, finding an appropriate tool for each data set would be another challenge for obtaining high quality bins. DASTool (Sieber, Probst et al. 2018) offers a solution for improving the quality of the bins. DASTool is a dereplication, aggregation and scoring automated tool that integrates several binning tool’s outputs and creates an optimized and non-redundant bin. To validate the performance of DASTool algorithm, DASTool was tested for simulated metagenomic data sets, environmental samples and metagenomic data with different complexity level. For simulated metagenomic data sets, the developers of DASTool used three different metagenomic data set created including low complexity sample (40 genomes), medium complexity (132 genomes), and high complexity (596 genomes). For environmental samples, metagenomic data from a high-CO2 cold-water geyser was used. To evaluate DASTool algorithm for different complexity metagenomic data sets, metagenomic shotgun sequencing data used from human microbiome, natural oil steeps and soil. These three groups of data sets were clustered using five different unsupervised binning tools (four hybrid tool ABAWACA 1.07 (https://github.com/CK7/abawaca), CONCOCT (Alneberg, Bjarnason et al. 2014), MaxBin 2 (Wu, Simmons et al. 2015), MetaBAT (Kang, Froula et al. 2015); and one nucleotide composition tool tetranucleotide ESOMs (Dick, Andersson et al. 2009)). Then, DASTool used to optimize the result of these individual binning tools. In all cases DASTool outperformed the result of individual binning tools by improving the quality of bins (low contamination and high completeness). Thus, in our pipeline (CuBi-MeAn) we used DASTool to enhance the quality of the bins generated with our selected binning tools. User may refer to DASTool webpage for more information (https://github.com/cmks/DAS_Tool).

##### 1.1.2. Downstream Data Assessment and Analysis

###### Quality Assessments

For quality assessment of the bins, CheckM software (Parks, Imelfort et al. 2015) was used to evaluate the bins generated by binning tools and DASTool. Quality of bins generated from metagenomic data are a major factor impacting the performance of the binning tools. In metagenomic assembled genomes, unlike the single isolate genome assembly, the genomes are recovered from a diverse group of microorganisms, therefore there is always the potential to introduce DNA fragments into the metagenomic assembled bins that are not actually belongs to. Identification and quantification of the universal single copy genes (USCG) present in the bins are one of the most common approaches to evaluate the quality of the MAGs. The quality of the MAGs is commonly assessed by calculating contamination and completeness. Completeness is the number of unique USCGs present within the bin. Conversely, contamination is the number of USCGs present in multiple copies, as only one copy should be present of each USCG per genome. Among available quality assessment tools, CheckM is a popular one which comprised of a set of tools for assessing the quality of genomes recovered metagenomic bins, metagenomes, new isolates genomes etc. It provides robust estimates of genome completeness and contamination by using sets of single copy and ubiquitous genes within a phylogenetic lineage. For more information user can review CheckM tool page (https://github.com/Ecogenomics/CheckM/wiki).

###### Taxonomical Classifications

Taxonomic classification of the bins was performed by using CheckM, PhyloPhlAn (Segata, Börnigen et al. 2013) and CAT/BAT (von Meijenfeldt, Arkhipova et al. 2019).

For taxonomical classifications CheckM relies on single copy marker genes that are specific to a genome’s lineage within a reference genome tree. CheckM uses 104 linage specific marker data sets taxonomical classification.

PhyloPhlAn uses the most conserved 400 proteins for extracting the phylogenetic signal. The marker gene identification step aims at first selecting the most relevant and the highest number of phylogenetic markers for the input sequences and then identifying them in the input sequences. The selection of the markers depends on the type of phylogeny considered and ranges from the 400 universal proteins to a variable number of core genes and species-specific genes.

CAT/BAT used the DIAMOND protein database (Buchfink, Xie et al. 2015) and Last Common Ancestors (LCA) for taxonomic classification. CAT/BAT algorithm involves gene calling, mapping of predicted open reading frames (ORFs) against the protein database, and voting-based classification of the entire contigs in the assembled genomes based on classification of the individual ORF(von Meijenfeldt, Arkhipova et al. 2019)..

To quantify the microbial community profile, bins mapped against the original metagenomic data to find out the alignment rate of contigs in the bins in the entire metagenomic data. In this study Bowtite2 software package (Langmead and Salzberg 2012) used for mapping the bins.

###### Functional Annotation

For functional annotation CuBi-MeAn used available online platforms KBase (Arkin, Cottingham et al. 2018) and RAST (Aziz, Bartels et al. 2008). These platforms which equipped with several annotation tools that users can choose, utilize and compare the results. Both of these platforms are equipped with SEED (Overbeek, Olson et al. 2014) which can provide a powerful tool for annotation. One of the advantages of SEED is that it updates constantly and has been linked to several other sources such as Swiss-Prot (Apweiler, Martin et al. 2010), GenBank (Benson, Cavanaugh et al. 2012), KEGG (Kanehisa, Goto et al. 2012) which offers user more options in dispose to annotate the bins in the same platform. Both of these platforms are equipped with other popular tools such as BLAST. In addition, KBase are also includes tools that are for metagenomic sequence analysis such as CheckM and assembly tools such as IDBA.

## 4. Results and Discussions

CuBi-MeAn was developed and applied to carry out a three-pronged approach to the analysis of metagenomic data from three different environmental engineering projects (TNT contaminated soil, EBPR reactor and algae-bacteria bioreactor). These three-pronged approaches are as follow:

Approach 1: To understand the dynamics of microbial community structure; Approach 2: To understand microbial function in the given environment; Approach 3: To explain how environmental factors affect the microbial communities. In this section, the performance of CuBi-MeAn in processing three different metagenomic datasets was evaluated. The information regarding the data types and other metrics are summarized in Table 2.1. The performance of CuBi-MeAn pipeline for abovementioned data sets will be discussed in this section.

**Table 2. 1:**
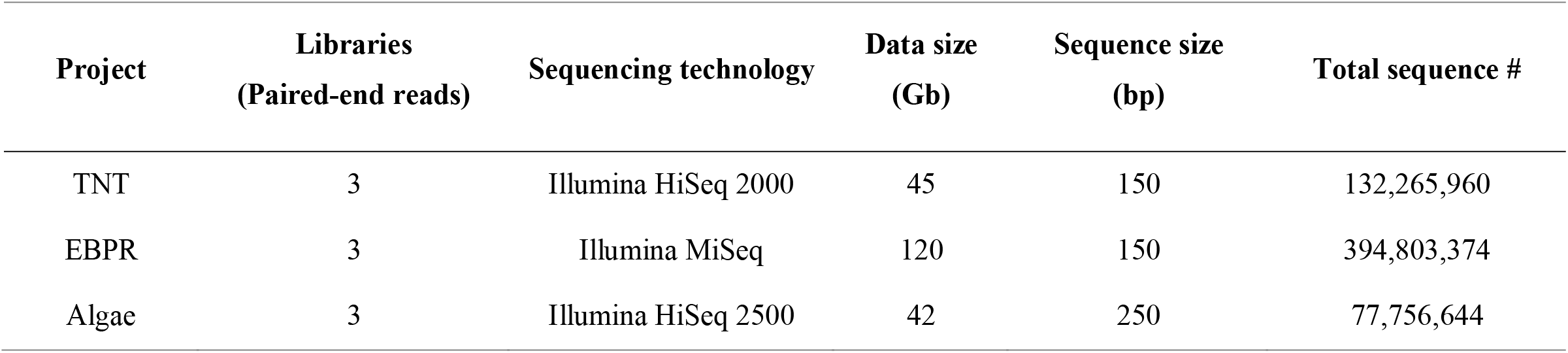
Information and certain metrics of metagenomic libraries tested by CuBi-MeAn pipeline.

The metagenomic data are generated from different origins. The first data sets, TNT was generated by collecting soil samples contaminated with an old TNT manufacturing site. The soil contains high concentrations of nitroaromatic compounds such as TNT, DNTs etc. The site was under aeration treatments by providing periodic tilling for 6 years. The second data set are EBPR data collected from aqueous sample. In this study bench-scale enhanced biological phosphorous removal (EBPR) reactors was designed and operated to study the organic phosphorous uptake from synthetic wastewater by certain microorganisms. The metagenomic samples in this study are from the EBPR reactor after 23 days of aerobic-anaerobic cycles. Finally, Algae data which also sequenced from aqueous samples are collected from an algae-bacteria bioreactor designed to enhance nitrogen removal from wastewater. As mentioned, and shown in Table 2.1, these metagenomic samples have different properties therefore we expect to have different performance from the pipeline which is used in this study.

### 1.1.3. Assembly

In this study three de-novo assembly tools Velvet, IDBA and Celera assembly were tested initially to evaluate the performance and computational requirements of these tools; Then, one assembly tool selected for downstream analysis. Assembly of the metagenomic data is one of the most computational resource intensive steps which requires high storage and RAM. This is a major constraint which needs to be considered for developing pipelines or simply processing metagenomic data sets. Therefore, in our study, four different subsample raw reads extracted from TNT data sets were tested to compare the performance of selected assemblers. These three reads were generated by mapping raw reads to selected reference genomes. The results of this test are summarized in Table 2.2.

**Table 2. 2:**
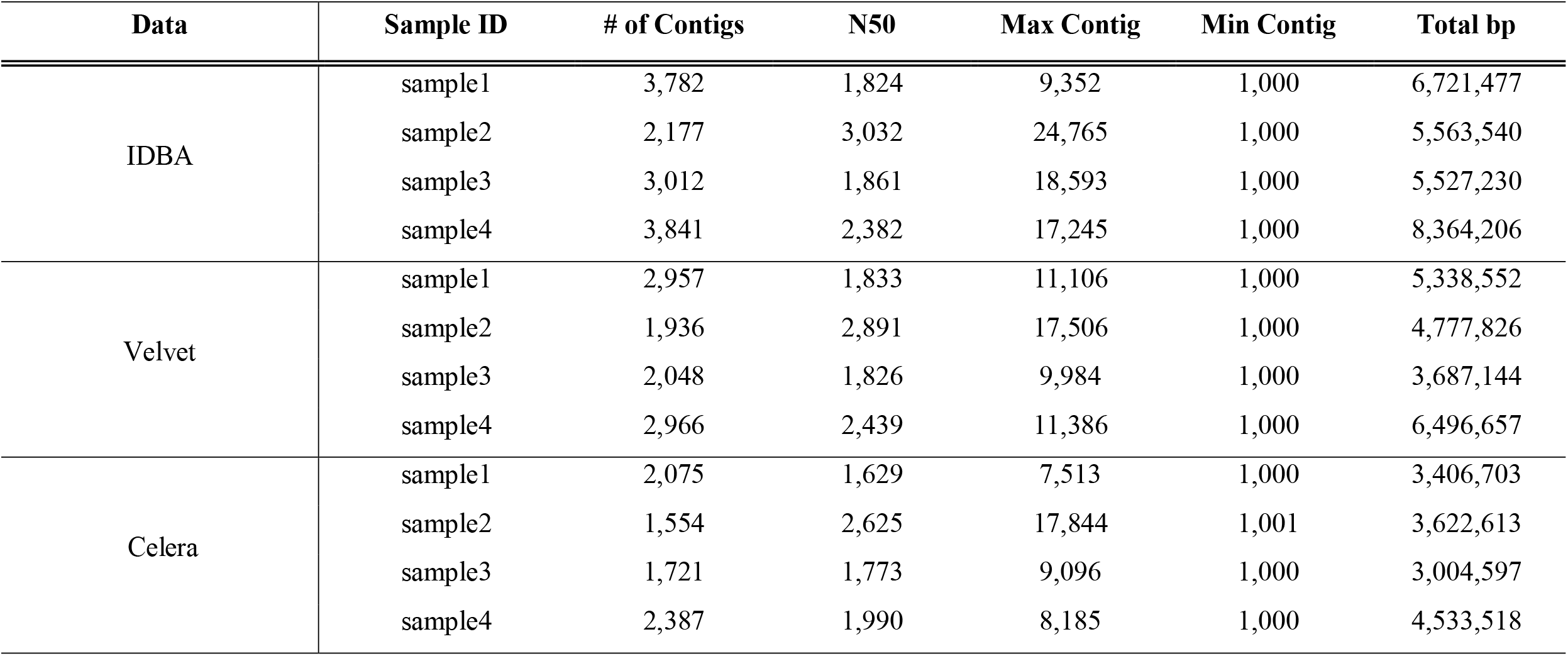
Assembly evaluations of IDBA, Velvet and Celera assembly tested for four different raw read samples.

Among these assemblers, IDBA had the better performance overall, with Velvet performing better than Celera. Also, Celera assembler was not specifically designed to handle metagenomic samples, while the two others had options to process metagenomic data. Therefore, for the next test in this study, we only compared the assemblies generated by IDBA and Velvet for the unfiltered EBPR and TNT datasets.

For both IDBA and Velvet, large “kmer” files are generated, which tabulates the number of occurrences for each fixed-length word of length k in a DNA data set. Generating the kmer file is an extremely time-consuming task and it usually produces an intermediate file that requires a large amount of storage to run properly. In our tests, TNT generated a ∼170 Gb and EBPR generated a ∼310 Gb kmer file. The next steps after generating kmer hash tables is contigs generation—the most RAM extensive task. For IDBA assembly, it used up almost 220Gb and 240Gb of RAM while it never went through for Velvet to complete the assembly. This has been also confirmed previously that IDBA is one of the least memory intensive assembly tools among other popular assembly tools (Abbas, Malluhi et al. 2014, van der Walt, van Goethem et al. 2017). Since, our disposable RAM for this project was 256 Gb, due to memory limitation IDBA was our option for metagenomic assembly. The running time for IDBA to be completed and generate final contigs file was almost 36 hours for TNT and 42 hours for EBPR data. It could have been possible to choose Velvet or other assembler such as MetaSPAdes (Nurk, Meleshko et al. 2017) or MEGAHIT (Li, Liu et al. 2015). if we had better computing/memory resources in dispose.

### 1.1.4. Raw data quality filtering

Quality filtering of the raw reads removes low quality reads that generate at the sequence reading step by sequencing facilities as non-determinized sequences or “N” (Figure 2.2). The low-quality reads with deteriorating quality reads are more observed towards the 3’-end, but they can be observed towards the 5’-end as well. These, incorrectly called bases negatively impact assembles, mapping, and downstream bioinformatics analysis (Young, Abaan et al. 2010). The assembled metagenomes from the assembly step are used as reference for clustering of metagenomic for binning. Shorter contigs size could cause inaccurate ambiguous binning due to low-complexity repetitive sequence (Chaisson and Pevzner 2008). Thus, in this study the quality filtering of the raw reads was tested on the assembly performance.

**Figure 2. 2:**
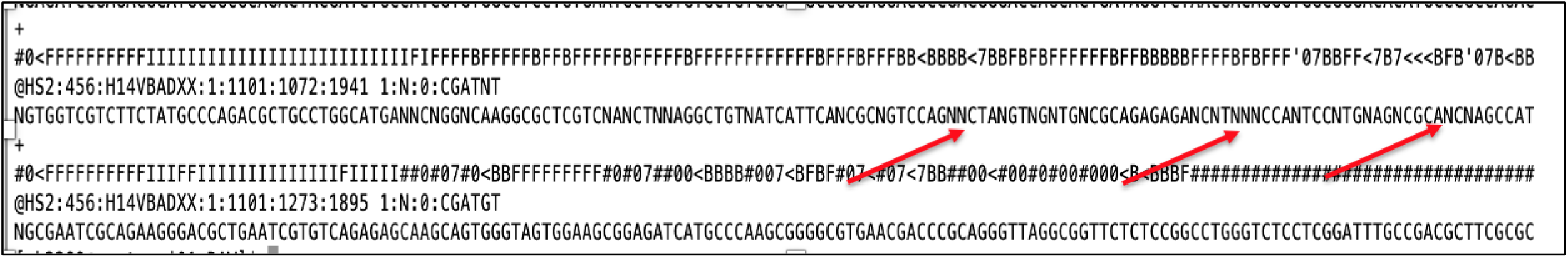
Fastq file raw reads from TNT soil sample generated by Illumina HiSeq 2000. The red arrows show non determined DNA nucleotides as “N”.

For this study initially no quality filtering of the raw metagenomic libraries was performed, which resulted in shorter and less completed contigs. Then, the metagenomic libraries were filtered to remove the low-quality sequences. In order to demonstrate the benefits of quality filtering the raw reads, we compared the summary statistics of the generated contigs of filtered and unfiltered reads when assembled by IDBA.

The results for TNT data are shown Table 2.3. Quality filtering of the raw reads by Sickle tool improved the quality of the assembled metagenome by increasing the average length (e.g., N50, N90 and N95) of contigs generated by assembly tool (Table 2.3).

**Table 2. 3:** Assembly evaluation for unfiltered assembled metagenomic reads (left) vs. filtered assembled metagenomic reads by Sickle tool (right) for TNT (a), EBPR (b) and Algae (c) metagenomic data.

| Assembly contigs unfiltered reads |  | Assembly contigs filtered reads |  |
| --- | --- | --- | --- |
| #Seqs | 37,302,779 | #Seqs | 4,109,944 |
| Min | 100 | Min | 100 |
| 1st Qu. | 118 | 1st Qu. | 193 |
| Median | 152 | Median | 240 |
| Mean | 201 | Mean | 364 |
| 3rd Qu. | 230 | 3rd Qu. | 338 |
| Max | 7,679 | Max | 128,391 |
| Total | 4,009,743,629 | Total | 1,497,876,890 |
| n50 | 192 | n50 | 386 |
| n90 | 116 | n90 | 188 |
| n95 | 109 | n95 | 173 |

**b.**
| Assembly contigs unfiltered reads |  | Assembly contigs filtered reads |  |
| --- | --- | --- | --- |
| #Seqs | 25,474,556 | #Seqs | 2,640,554 |
| Min | 100 | Min | 100 |
| 1st Qu. | 116 | 1st Qu. | 204 |
| Median | 174 | Median | 280 |
| Mean | 228 | Mean | 579 |
| 3rd Qu. | 350 | 3rd Qu. | 527 |
| Max | 17,872 | Max | 294,096 |
| Total | 2,040,146,547 | Total | 1,530,582,685 |
| n50 | 239 | n50 | 923 |
| n90 | 173 | n90 | 223 |

| n95 | 158 | n95 | 188 |
| --- | --- | --- | --- |
| **c.** |  |  |  |
| Assembly contigs unfiltered reads |  | Assembly contigs filtered reads |  |
| #Seqs | 33,155,430 | #Seqs | 3,607,782 |
| Min | 100 | Min | 100 |
| 1st Qu. | 113 | 1st Qu. | 141 |
| Median | 142 | Median | 206 |
| Mean | 159 | Mean | 399 |
| 3rd Qu. | 171 | 3rd Qu. | 320 |
| Max | 1,783 | Max | 149,090 |
| Total | 4,784,033,344 | Total | 184,281,121 |
| n50 | 156 | n50 | 600 |
| n90 | 117 | n90 | 174 |
| n95 | 112 | n95 | 123 |

### 1.1.5. Binning and bin refinement

As discussed in the first chapter we adopted a genome-centric approaches to study the systems used in this study; Thus, we processed our metagenomic data sets using binning approaches. Binning of metagenomic reads approximates the functions and taxonomy of the assigned genomes, while bypasses the challenges of full genome assembly. (Ribeca and Valiente 2011, Imelfort, Parks et al. 2014). Metagenomic assembled genomes (MAGs) or bins includes core genes of closely related taxa that has common genes and functions and at the same time pan-genes contains genes that are variably present in the bins (Tettelin, Masignani et al. 2005). Pan-genes have specific and specialized functions and adaptations of divergent taxonomical units. Therefore, binning could appropriately address challenges of genome centric approaches of diverse metagenomic samples.

Here, five different binning tools were used to process our three subject metagenomic datasets. These tools used different approaches (i.e. different algorithms) for processing metagenomic data, which results in low quality and incomplete bins for some data sets and better performance for other data sets using the same tool. Thus, finding an appropriate tool for each data set would be another challenge for obtaining high quality bins. DASTool (Sieber, Probst et al. 2018) offers a solution for improving the quality of the bins.

The selected binning tools in this study were BinSanity, MaxBin2, MetaBAT, COCACOLA and CONCOCT which are all hybrid clustering methods that use both kmer frequency and samples’ co-abundance. Then, DASTool was used as a refinement tool to select best quality contigs among different binning tools outputs.

In this study, the quality of the bins was evaluated by CheckM. As discussed, earlier CheckM is an automated tool that uses a broader range of marker genes for quality assessments and taxonomical classification. CheckM offers several options that generates results to evaluate MAGs. Completeness and contaminations are two factors that are extensively used to assess the quality of MAGs. Contamination is the “false positive”, while completeness is “true positive” from single copy marker genes in the given MAGs. In our study, we also used mostly these two parameters for quality assessment of our bins. The CheckM results indicate that the quality of the bins generated by DASTool outperformed individual binning methods (Figure 2.3). However, there are some variation in quality of the bins generated by DASTool among different data sets (Figure 2.4). The results suggested that the quality of the DASTool bins depended upon the quality of the bins generated by individual binning tools. This could be explained by the approaches and algorithms of applied by DASTool to generate the bins. It means that DASTool is not an assembly or clustering tool, instead it generates the new bins by evaluation and aggregation of the best contigs from bins generated by other binning tools.

**Figure 2. 3:**
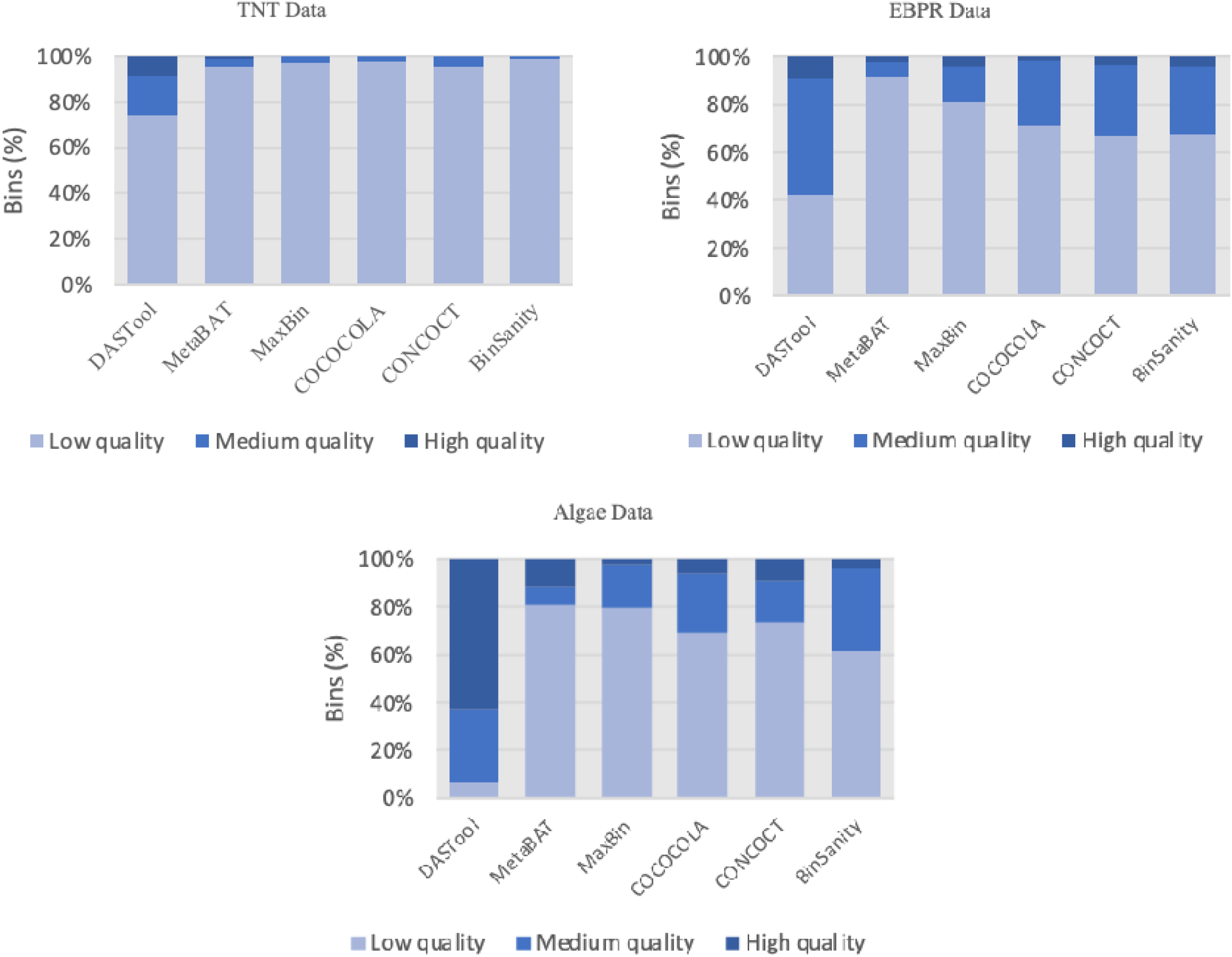
Quality comparison of bins generated by MetaBAT, MaxBin2, COCACOLA, CONCOCT, BinSanity and DASTool.

**Figure 2. 4:**
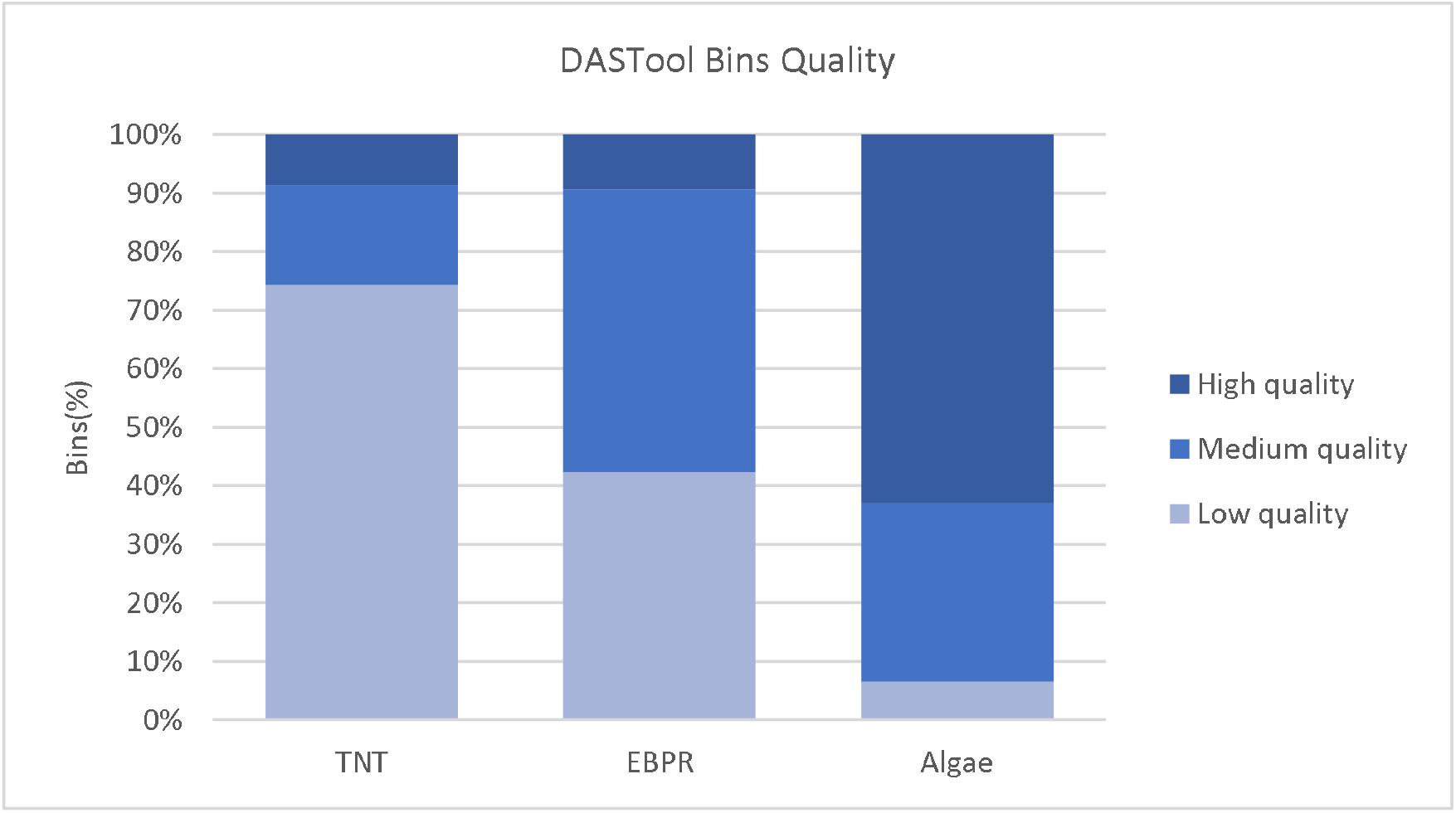
Bin quality comparison of three metagenomic data sets TNT, EBPR and Algae generated by DASTool.

Also, data source, coverage, reads length and technology used affect the quality of the final results. Among our data sets “Algae data” generates the best quality bins with less contaminations and more completeness (Figures 2.3 and 2.4). This could be explained by the fact that raw Algae data has the longest reads among others (Table 2.1). Longer reads would generate longer contigs which can be aligned and mapped with less ambiguity (Chaisson and Pevzner 2008). In addition, the origin of the sample (soil vs water, etc) could impact the quality of the bins. For example, TNT data is from the soil samples which inherently are very diverse, compared to aqueous samples, making clustering more difficult. Therefore, we have the least high and medium quality bins in TNT samples comparing to other data sets which are from water samples.

### 1.1.6. Bin analyses

Bins generated by DASTool were used for downstream analysis for data interpretation and addressing specific project research questions. CuBi-MeAn pipeline uses DASTool bins for: i. Taxonomical classification; ii. Functional annotation; And iii. Quantification of the microbial profile in the given system.

#### Taxonomical Classification

Taxonomical classification of the bins was performed with three different tools. CheckM uses linage specific marker sets for classification of the bins; PhyloPhlAn uses the most conserved 400 proteins for extracting the phylogenetic signal; And, CAT/BAT uses DAMOND protein databases and Last Common Ancestors (LCA) for taxonomic classification.

Different bins quality, different samples from different sources, and different classification approaches could be the reasons for different classifications among and within samples. Table 2.4 compares CheckM, PhyloPhlAn and CAT/BAT taxonomical classifications of the TNT bins as an example. Since, there are different approaches and databases used for taxonomical classifications there are not any preference or advantages of these tools over others. CheckM tree could be generated simply by adding a single script to CheckM data processing steps used in this pipeline for quality assessments of the bins. Instead, PhyloPhlAn needs more steps to generate phylogenetic trees. CAT/BAT also used a different approach as mentioned earlier and has a few simple steps which makes it an easier and faster tool compared to PhyloPhlAn. Overall, the users can use these classification tools or add new tools and choose the best of the results from the bins’ classification.

**Table 2. 4:**
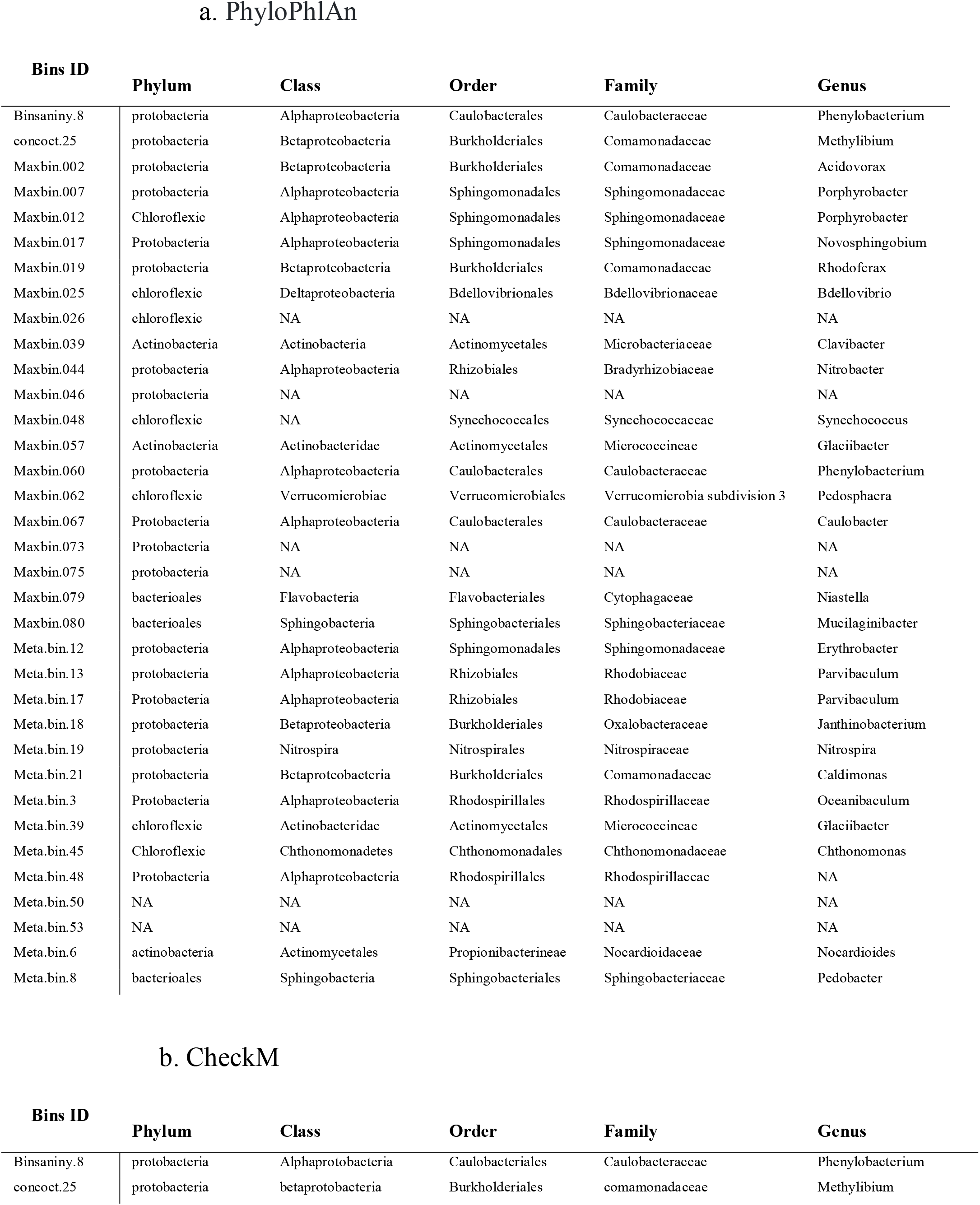
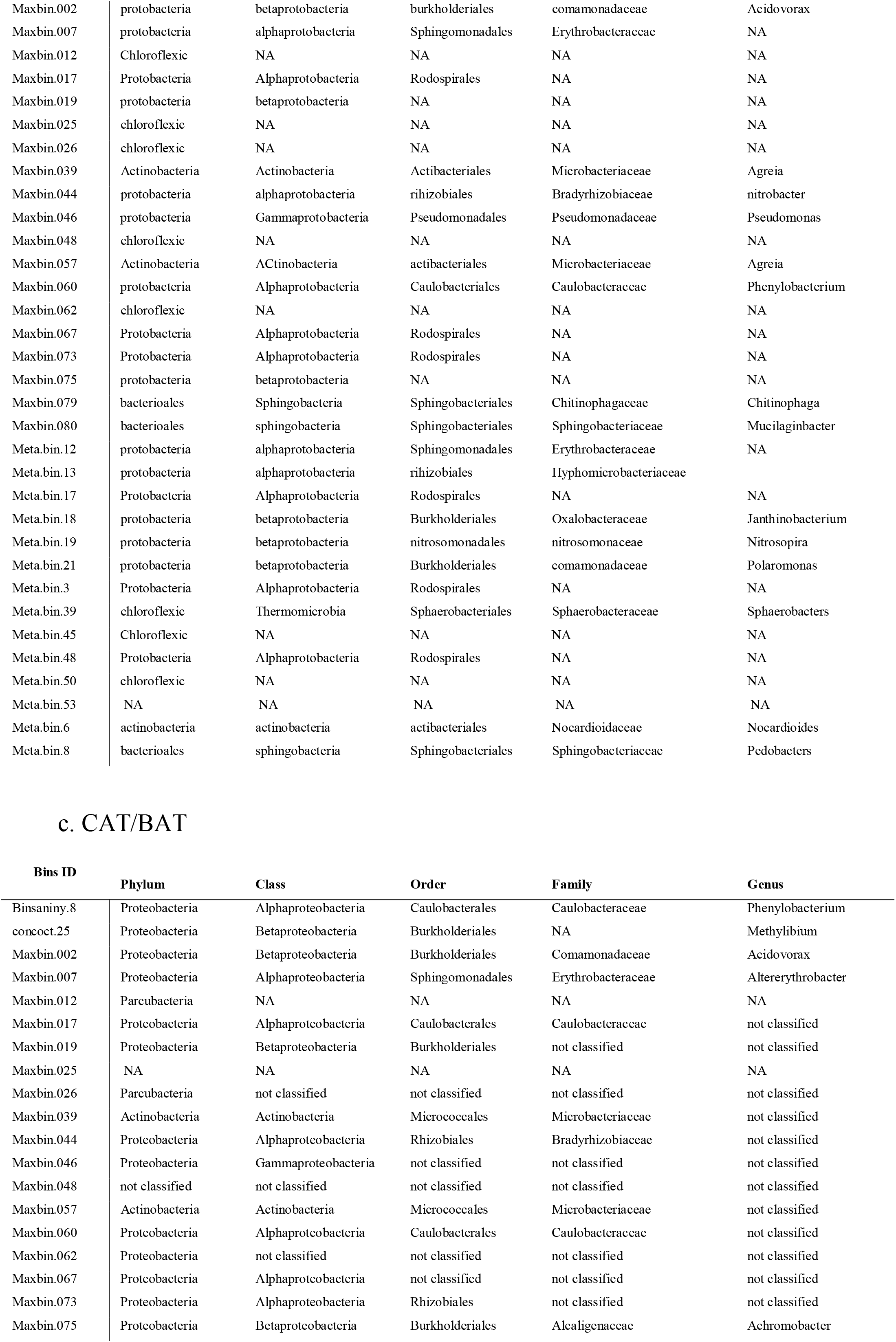
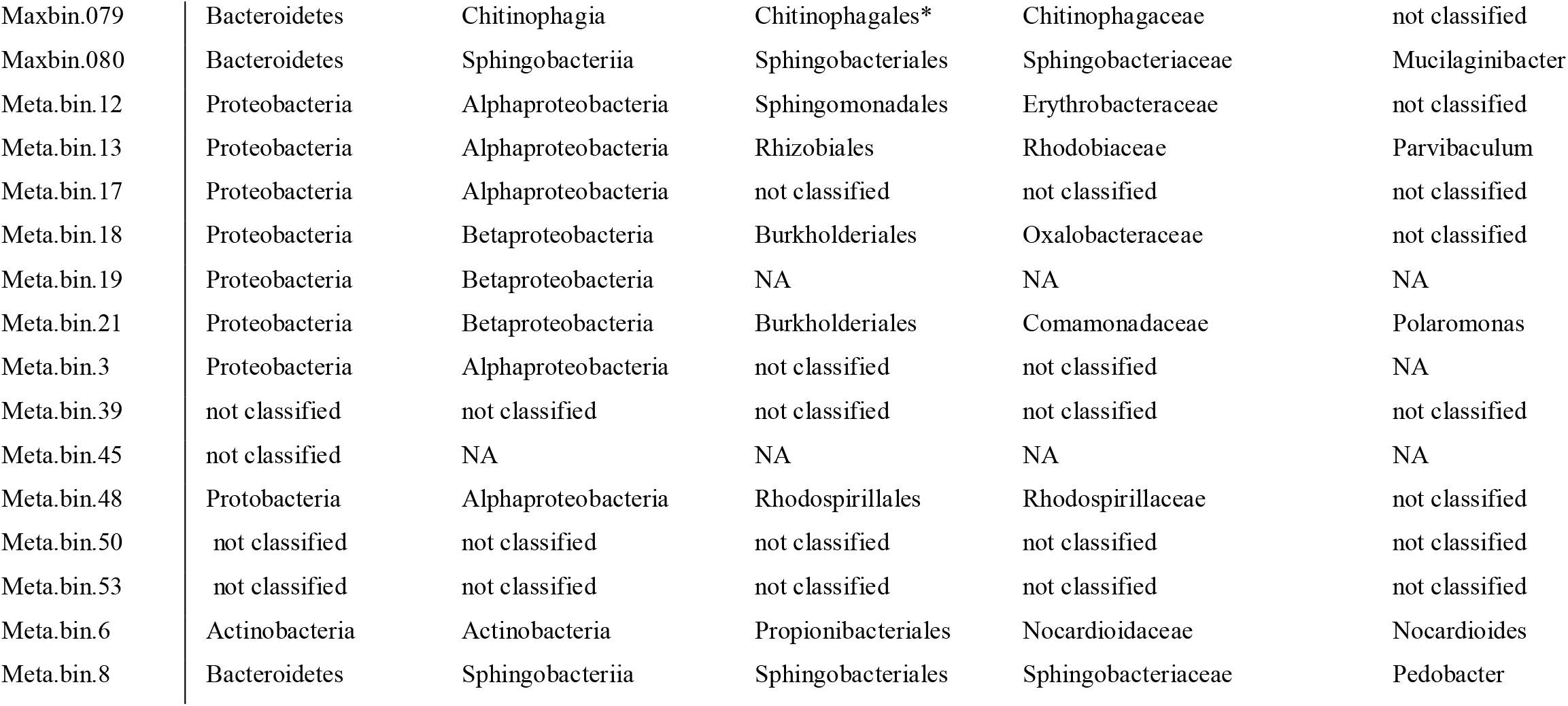
Taxonomical classifications of TNT bins by a. PhyloPhlAn, b. CheckM and c. CAT/BAT

#### Quantification of the microbial profile

To find out the microbial community profile in addition to taxonomical classifications their abundance is also investigated in this study. The abundance of certain groups of microorganisms in a system could explain why those group are more successful in that system in the given period of time, how they interact with chemical and biological composition of the environment, etc. In this study contigs in the MAGs were mapped to the metagenomic raw reads to evaluate the alignment rate of the contigs in the bins. The overall microbial community composition profile could explain the dynamic of our systems. For example, Figure 2.5 shows microbial community profile of Algae reactors. The results show strong presence of the predatory microorganisms in the system which plays an important role in the dynamics and community composition of the reactors.

**Figure 2. 5:**
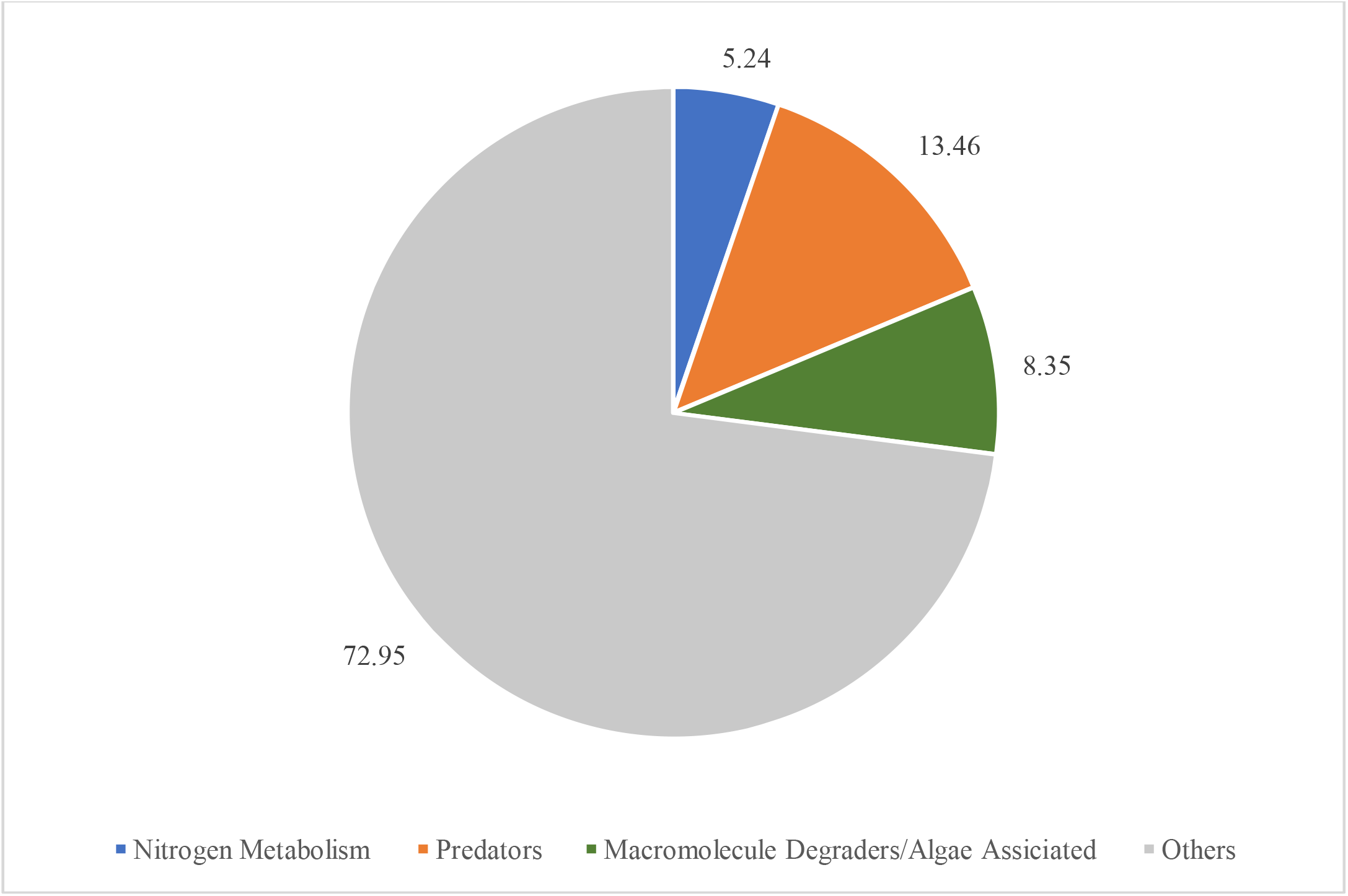
Microbial community profile in Algae reactor.

#### Functional annotation

For functional annotation of the bins generated by DASTool, CuBi-MeAn utilize two different online platforms Kbase and RAST. For three metagenomic datasets in this project SEED, GeneBank, UniProt, BLAST, etc. were used to identify the key genes in each system, construct metabolic pathways or identify and investigate other genes that their functions decode project specific research questions. Figure 2.6 demonstrates TNT complete degradation pathway of using these annotation tools. In this study certain genes were not detected in SEED database; therefore, other alternative such as alignment of those genes against bins by NCBI BLAST tool, was used for investigation.

**Figure 2. 6:**
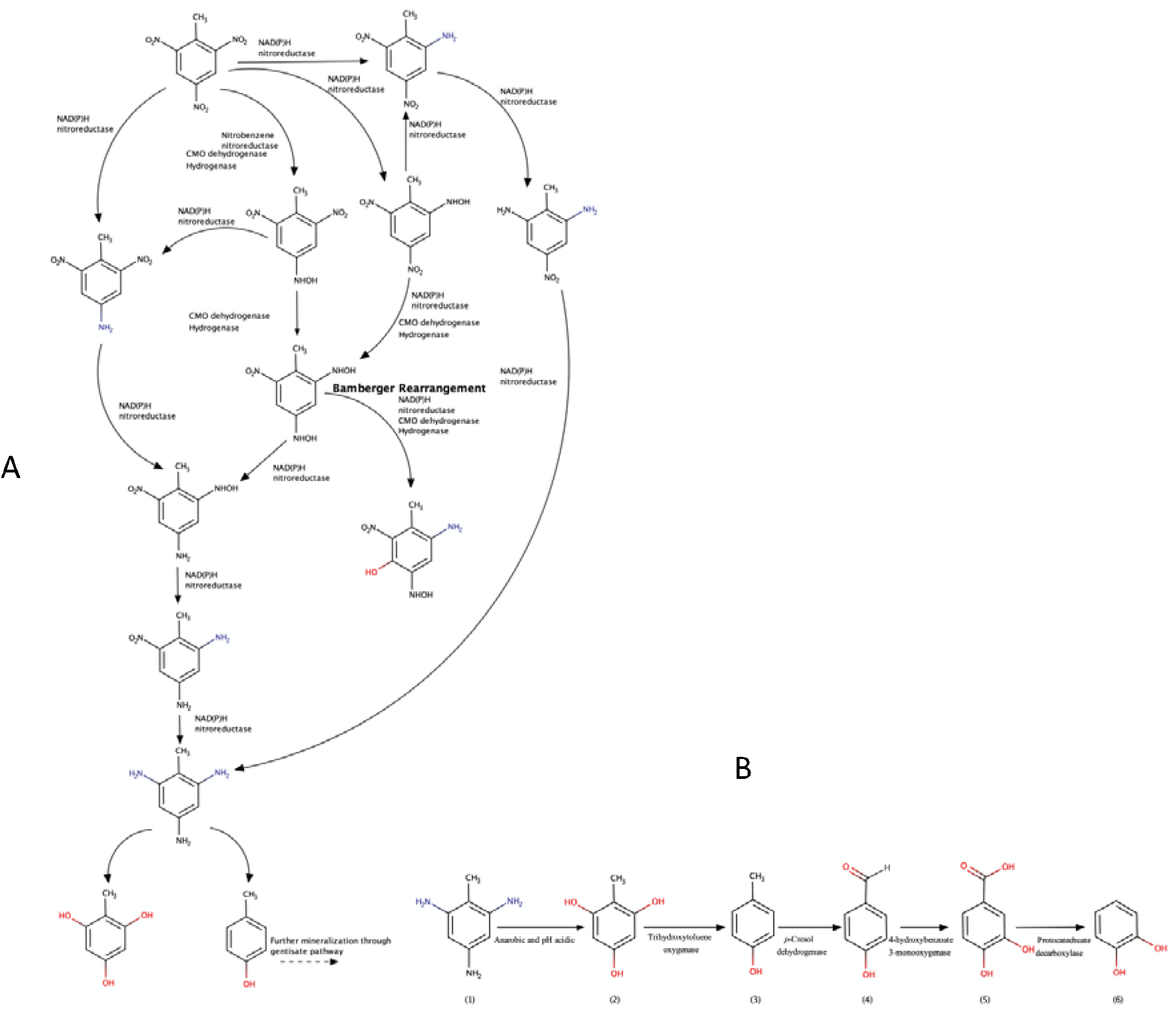
TNT degradation pathway constructed by using CuBi-MeAn In addition, online genome data bases such as GeneBank used as benchmark to compare.

and investigate our constructed MAGs. For example, in Algae project there are certain bacterial guilds in the reactor were suggested to have some sorts of defense mechanism against the predation. We hypothesized that could be the reason why those groups of bacteria are more abundant in our reactors. Previous studies suggested that these defense mechanisms in the MAGs were certain genes and some specific DNA structures known as clustered regularly interspaced short palindromic repeats (CRISPRs) elements which could be an indication of the defense mechanisms. However, these defense elements were not detected in SEED database using KBase and RAST platforms. Instead, the genes of the defense mechanism genes were obtained from the GeneBank database and BLAST tool used to test the gene presence in our bins. BLAST tools aligned these genes to our bins with high scores. For CRISPRs detection we also used another tool that was designed specifically for this purpose (Edgar 2007). The gene annotation of the bins helps to find out to investigate and understand the dynamics of the entire system for TNT, EBPR and Alga projects. These results will be discussed extensively in projects specific sections in the following sections.

## 5. Conclusions

In this study a customized pipeline was developed to process and analyze the metagenomic libraries. CuBi-MeAn pipeline was used for investigation of microbial community profile and functional annotations. This study was demonstrated how a genome centric approach could explain the functions of environmental systems and answer questions underlying the dynamics of the systems by using this pipeline. In this study, CuBi-MeAn pipeline clusters the metagenomic reads to approximate the genome of microorganisms exist in that system for downstream analysis. In addition, to the clustering of the metagenomic reads using this pipeline, this study showed how the selected tools in CuBi-MeAn pipeline, improved the quality of the raw data which enhanced the metagenomic assembly, generate bins, and downstream analysis. By designing and developing a flexible and customized pipeline, this study showed how to process large metagenomic data sets with limited resources.

Our proof-of-concept can be applied to process similar metagenomic datasets from short read metagenomic sequences of environmental samples. The user of CuBi-MeAn would be able to update, customize, replace, or skip specific software or steps. Since this pipeline is comprised of several metagenomics tools, they could be performed sequentially on different platforms and machines based on their available resources.

Despite successful demonstration of CuBi-MeAn, still there is room for further development of this pipeline. Thereby, the authors plan to test the performance of CuBi-MeAn with a wide variety of datasets such as human microbiome and using different sequences technologies such as PacBio sequencer that generate long reads.

## References

Abbas, M. M., Q. M. Malluhi and P. Balakrishnan (2014). “Assessment of de novo assemblers for draft genomes: a case study with fungal genomes.” BMC genomics 15 Suppl 9(Suppl 9): S10–S10.

Albertsen, M., P. Hugenholtz, A. Skarshewski, K. L. Nielsen, G. W. Tyson and P. H. Nielsen (2013). “Genome sequences of rare, uncultured bacteria obtained by differential coverage binning of multiple metagenomes.” Nature biotechnology 31(6): 533–538.

Almeida, A., A. L. Mitchell, M. Boland, S. C. Forster, G. B. Gloor, A. Tarkowska, T. D. Lawley and R. D. Finn (2019). “A new genomic blueprint of the human gut microbiota.” Nature 568(7753): 499–504.

Alneberg, J., B. S. Bjarnason, I. De Bruijn, M. Schirmer, J. Quick, U. Z. Ijaz, L. Lahti, N. J. Loman, A. F. Andersson and C. Quince (2014). “Binning metagenomic contigs by coverage and composition.” Nature methods 11(11): 1144.

Alneberg, J., B. S. Bjarnason, I. de Bruijn, M. Schirmer, J. Quick, U. Z. Ijaz, L. Lahti, N. J. Loman, A. F. Andersson and C. Quince (2014). “Binning metagenomic contigs by coverage and composition.” Nature Methods 11(11): 1144–1146.

Anantharaman, K., C. T. Brown, L. A. Hug, I. Sharon, C. J. Castelle, A. J. Probst, B. C. Thomas, A. Singh, M. J. Wilkins, U. Karaoz, E. L. Brodie, K. H. Williams, S. S. Hubbard and J. F. Banfield (2016). “Thousands of microbial genomes shed light on interconnected biogeochemical processes in an aquifer system.” Nature Communications 7(1): 13219.

Apweiler, R., M. J. Martin, C. O’Donovan, M. Magrane, Y. Alam-Faruque, R. Antunes, D. Barrell, B. Bely, M. Bingley and D. Binns (2010). “The universal protein resource (UniProt) in 2010.” Nucleic Acids Research 38: D142–D148.

Arkin, A. P., R. W. Cottingham, C. S. Henry, N. L. Harris, R. L. Stevens, S. Maslov, P. Dehal, D. Ware, F. Perez, S. Canon, M. W. Sneddon, M. L. Henderson, W. J. Riehl, D. Murphy-Olson, S. Y. Chan, R. T. Kamimura, S. Kumari, M. M. Drake, T. S. Brettin, E. M. Glass, D. Chivian, D. Gunter, D. J. Weston, B. H. Allen, J. Baumohl, A. A. Best, B. Bowen, S. E. Brenner, C. C. Bun, J.-M. Chandonia, J.-M. Chia, R. Colasanti, N. Conrad, J. J. Davis, B. H. Davison, M. DeJongh, S. Devoid, E. Dietrich, I. Dubchak, J. N. Edirisinghe, G. Fang, J. P. Faria, P. M. Frybarger, W. Gerlach, M. Gerstein, A. Greiner, J. Gurtowski, H. L. Haun, F. He, R. Jain, M. P. Joachimiak, K. P. Keegan, S. Kondo, V. Kumar, M. L. Land, F. Meyer, M. Mills, P. S. Novichkov, T. Oh, G. J. Olsen, R. Olson, B. Parrello, S. Pasternak, E. Pearson, S. S. Poon, G. A. Price, S. Ramakrishnan, P. Ranjan, P. C. Ronald, M. C. Schatz, S. M. D. Seaver, M. Shukla, R. A. Sutormin, M. H. Syed, J. Thomason, N. L. Tintle, D. Wang, F. Xia, H. Yoo, S. Yoo and D. Yu (2018). “KBase: The United States Department of Energy Systems Biology Knowledgebase.” Nature Biotechnology 36(7): 566–569.

Aziz, R. K., D. Bartels, A. A. Best, M. DeJongh, T. Disz, R. A. Edwards, K. Formsma, S. Gerdes, E. M. Glass and M. Kubal (2008). “The RAST Server: rapid annotations using subsystems technology.” BMC genomics 9(1): 75.

Bashiardes, S., G. Zilberman-Schapira and E. Elinav (2016). “Use of metatranscriptomics in microbiome research.” Bioinformatics and biology insights 10: BBI. S34610.

Benson, D. A., M. Cavanaugh, K. Clark, I. Karsch-Mizrachi, D. J. Lipman, J. Ostell and E. W. Sayers (2012). “GenBank.” Nucleic acids research 41(D1): D36-D42.

Brown, C. T., L. A. Hug, B. C. Thomas, I. Sharon, C. J. Castelle, A. Singh, M. J. Wilkins, K. C. Wrighton, K. H. Williams and J. F. Banfield (2015). “Unusual biology across a group comprising more than 15% of domain Bacteria.” Nature 523(7559): 208–211.

Buchfink, B., C. Xie and D. H. Huson (2015). “Fast and sensitive protein alignment using DIAMOND.” Nature Methods 12(1): 59–60.

Castelle, C. J., K. C. Wrighton, B. C. Thomas, L. A. Hug, C. T. Brown, M. J. Wilkins, K. R. Frischkorn, S. G. Tringe, A. Singh and L. M. Markillie (2015). “Genomic expansion of domain archaea highlights roles for organisms from new phyla in anaerobic carbon cycling.” Current biology 25(6): 690–701.

Chaisson, M. J. and P. A. Pevzner (2008). “Short read fragment assembly of bacterial genomes.” Genome research 18(2): 324–330.

Clarke, E. L., L. J. Taylor, C. Zhao, A. Connell, J.-J. Lee, B. Fett, F. D. Bushman and K. Bittinger (2019). “Sunbeam: an extensible pipeline for analyzing metagenomic sequencing experiments.” Microbiome 7(1): 1–13.

Cole, S. T., R. Brosch, J. Parkhill, T. Garnier, C. Churcher, D. Harris, S. V. Gordon, K. Eiglmeier, S. Gas, C. E. Barry, F. Tekaia, K. Badcock, D. Basham, D. Brown, T. Chillingworth, R. Connor, R. Davies, K. Devlin, T. Feltwell, S. Gentles, N. Hamlin, S. Holroyd, T. Hornsby, K. Jagels, A. Krogh, J. McLean, S. Moule, L. Murphy, K. Oliver, J. Osborne, M. A. Quail, M. A. Rajandream, J. Rogers, S. Rutter, K. Seeger, J. Skelton, R. Squares, S. Squares, J. E. Sulston, K. Taylor, S. Whitehead and B. G. Barrell (1998). “Deciphering the biology of Mycobacterium tuberculosis from the complete genome sequence.” Nature 393(6685): 537–544.

Cotillard, A., S. P. Kennedy, L. C. Kong, E. Prifti, N. Pons, E. Le Chatelier, M. Almeida, B. Quinquis, F. Levenez, N. Galleron, S. Gougis, S. Rizkalla, J.-M. Batto, P. Renault, J. Doré, J.-D. Zucker, K. Clément, S. D. Ehrlich, H. Blottière, M. Leclerc, C. Juste, T. de Wouters, P. Lepage, C. Fouqueray, A. Basdevant, C. Henegar, C. Godard, M. Fondacci, A. Rohia, F. Hajduch, J. Weissenbach, E. Pelletier, D. Le Paslier, J.-P. Gauchi, J.-F. Gibrat, V. Loux, W. Carré, E. Maguin, M. van de Guchte, A. Jamet, F. Boumezbeur, S. Layec, A. N. R. M. consortium and A. N. R. M. c. members (2013). “Dietary intervention impact on gut microbial gene richness.” Nature 500(7464): 585–588.

Dawkins, R. (2016). The selfish gene, Oxford university press.

Dick, G. J., A. F. Andersson, B. J. Baker, S. L. Simmons, B. C. Thomas, A. P. Yelton and J. F. Banfield (2009). “Community-wide analysis of microbial genome sequence signatures.” Genome biology 10(8): 1–16.

Edgar, R. C. (2007). “PILER-CR: fast and accurate identification of CRISPR repeats.” BMC bioinformatics 8(1): 18.

Gohl, D. M., P. Vangay, J. Garbe, A. MacLean, A. Hauge, A. Becker, T. J. Gould, J. B. Clayton, T. J. Johnson and R. Hunter (2016). “Systematic improvement of amplicon marker gene methods for increased accuracy in microbiome studies.” Nature biotechnology 34(9): 942–949.

Graham, E. D., J. F. Heidelberg and B. J. Tully (2017). “BinSanity: unsupervised clustering of environmental microbial assemblies using coverage and affinity propagation.” PeerJ 5: e3035.

Guénet, J. L. (2005). “The mouse genome.” Genome research 15(12): 1729–1740.

Heng, H. H. (2009). “The genome_Jcentric concept: resynthesis of evolutionary theory.” Bioessays 31(5): 512–525.

Heyer, R., K. Schallert, A. Büdel, R. Zoun, S. Dorl, A. Behne, F. Kohrs, S. Püttker, C. Siewert, T. Muth, G. Saake, U. Reichl and D. Benndorf (2019). “A Robust and Universal Metaproteomics Workflow for Research Studies and Routine Diagnostics Within 24 h Using Phenol Extraction, FASP Digest, and the MetaProteomeAnalyzer.” Frontiers in Microbiology 10(1883).

Hiergeist, A., J. Gläsner, U. Reischl and A. Gessner (2015). “Analyses of intestinal microbiota: culture versus sequencing.” ILAR journal 56(2): 228–240.

Hodkinson, B. P. and E. A. Grice (2015). “Next-generation sequencing: a review of technologies and tools for wound microbiome research.” Advances in wound care 4(1): 50–58.

Holmes, A. J., M. R. Gillings, B. S. Nield, B. C. Mabbutt, K. H. Nevalainen and H. Stokes (2003). “The gene cassette metagenome is a basic resource for bacterial genome evolution.” Environmental microbiology 5(5): 383–394.

Imelfort, M., D. Parks, B. J. Woodcroft, P. Dennis, P. Hugenholtz and G. W. Tyson (2014). “GroopM: an automated tool for the recovery of population genomes from related metagenomes.” PeerJ 2: e603.

Jaenicke, S., C. Ander, T. Bekel, R. Bisdorf, M. Dröge, K.-H. Gartemann, S. Jünemann, O. Kaiser, L. Krause and F. Tille (2011). “Comparative and joint analysis of two metagenomic datasets from a biogas fermenter obtained by 454-pyrosequencing.” PloS one 6(1): e14519.

Jain, M., H. E. Olsen, B. Paten and M. Akeson (2016). “The Oxford Nanopore MinION: delivery of nanopore sequencing to the genomics community.” Genome biology 17(1): 239.

Joshi, N. and J. Fass (2011). Sickle: A sliding-window, adaptive, quality-based trimming tool for FastQ files (Version 1.33)[Software].

Jovel, J., J. Patterson, W. Wang, N. Hotte, S. O’Keefe, T. Mitchel, T. Perry, D. Kao, A. L. Mason and K. L. Madsen (2016). “Characterization of the gut microbiome using 16S or shotgun metagenomics.” Frontiers in microbiology 7: 459.

Juengst, E. and J. Huss (2009). “From metagenomics to the metagenome: Conceptual change and the rhetoric of translational genomic research.” Genomics, Society and Policy 5(3): 1.

Juengst, E. T. (2009). Metagenomic metaphors: New images of the human from ‘translational’genomic research. New Visions of Nature, Springer: 129–145.

Kanehisa, M., S. Goto, Y. Sato, M. Furumichi and M. Tanabe (2012). “KEGG for integration and interpretation of large-scale molecular data sets.” Nucleic acids research 40(D1): D109–D114.

Kang, D. D., J. Froula, R. Egan and Z. Wang (2015). “MetaBAT, an efficient tool for accurately reconstructing single genomes from complex microbial communities.” PeerJ 3: e1165.

Kougias, P. G., S. Campanaro, L. Treu, P. Tsapekos, A. Armani and I. Angelidaki (2018). “Spatial Distribution and Diverse Metabolic Functions of Lignocellulose-Degrading Uncultured Bacteria as Revealed by Genome-Centric Metagenomics.” Applied and Environmental Microbiology 84(18): e01244–01218.

Langmead, B. and S. L. Salzberg (2012). “Fast gapped-read alignment with Bowtie 2.” Nature Methods 9(4): 357–359.

Le Chatelier, E., T. Nielsen, J. Qin, E. Prifti, F. Hildebrand, G. Falony, M. Almeida, M. Arumugam, J.-M. Batto, S. Kennedy, P. Leonard, J. Li, K. Burgdorf, N. Grarup, T. Jørgensen, I. Brandslund, H. B. Nielsen, A. S. Juncker, M. Bertalan, F. Levenez, N. Pons, S. Rasmussen, S. Sunagawa, J. Tap, S. Tims, E. G. Zoetendal, S. Brunak, K. Clément, J. Doré, M. Kleerebezem, K. Kristiansen, P. Renault, T. Sicheritz-Ponten, W. M. de Vos, J.-D. Zucker, J. Raes, T. Hansen, E. Guedon, C. Delorme, S. Layec, G. Khaci, M. van de Guchte, G. Vandemeulebrouck, A. Jamet, R. Dervyn, N. Sanchez, E. Maguin, F. Haimet, Y. Winogradski, A. Cultrone, M. Leclerc, C. Juste, H. Blottière, E. Pelletier, D. LePaslier, F. Artiguenave, T. Bruls, J. Weissenbach, K. Turner, J. Parkhill, M. Antolin, C. Manichanh, F. Casellas, N. Boruel, E. Varela, A. Torrejon, F. Guarner, G. Denariaz, M. Derrien, J. E. T. van Hylckama Vlieg, P. Veiga, R. Oozeer, J. Knol, M. Rescigno, C. Brechot, C. M’Rini, A. Mérieux, T. Yamada, P. Bork, J. Wang, S. D. Ehrlich, O. Pedersen and H. I. T. c. Meta (2013). “Richness of human gut microbiome correlates with metabolic markers.” Nature 500(7464): 541–546.

Li, D., C. M. Liu, R. Luo, K. Sadakane and T. W. Lam (2015). “MEGAHIT: an ultra-fast single-node solution for large and complex metagenomics assembly via succinct de Bruijn graph.” Bioinformatics 31(10): 1674–1676.

Lu, Y. Y., T. Chen, J. A. Fuhrman and F. Sun (2017). “COCACOLA: binning metagenomic contigs using sequence COmposition, read CoverAge, CO-alignment and paired-end read LinkAge.” Bioinformatics 33(6): 791–798.

Mande, S. S., M. H. Mohammed and T. S. Ghosh (2012). “Classification of metagenomic sequences: methods and challenges.” Briefings in bioinformatics 13(6): 669–681.

Mardis, E. R. (2008). “The impact of next-generation sequencing technology on genetics.” Trends in genetics 24(3): 133–141.

Mendes, L. W., L. P. P. Braga, A. A. Navarrete, D. Souza, G. G. Silva and S. M. Tsai (2017). “Using metagenomics to connect microbial community biodiversity and functions.” Curr Issues Mol Biol 24: 103–118.

Mohammed, M. H., T. S. Ghosh, N. K. Singh and S. S. Mande (2011). “SPHINX—an algorithm for taxonomic binning of metagenomic sequences.” Bioinformatics 27(1): 22–30.

Myers, E. W., G. G. Sutton, A. L. Delcher, I. M. Dew, D. P. Fasulo, M. J. Flanigan, S. A. Kravitz, C. M. Mobarry, K. H. Reinert, K. A. Remington, E. L. Anson, R. A. Bolanos, H. H. Chou, C. M. Jordan, A. L. Halpern, S. Lonardi, E. M. Beasley, R. C. Brandon, L. Chen, P. J. Dunn, Z. Lai, Y. Liang, D. R. Nusskern, M. Zhan, Q. Zhang, X. Zheng, G. M. Rubin, M. D. Adams and J. C. Venter (2000). “A whole-genome assembly of Drosophila.” Science 287(5461): 2196–2204.

Namiki, T., T. Hachiya, H. Tanaka and Y. Sakakibara (2012). “MetaVelvet: an extension of Velvet assembler to de novo metagenome assembly from short sequence reads.” Nucleic Acids Res 40(20): e155.

Nielsen, H. B., M. Almeida, A. S. Juncker, S. Rasmussen, J. Li, S. Sunagawa, D. R. Plichta, L. Gautier, A. G. Pedersen, E. Le Chatelier, E. Pelletier, I. Bonde, T. Nielsen, C. Manichanh, M. Arumugam, J.-M. Batto, M. B. Quintanilha dos Santos, N. Blom, N. Borruel, K. S. Burgdorf, F. Boumezbeur, F. Casellas, J. Doré, P. Dworzynski, F. Guarner, T. Hansen, F. Hildebrand, R. S. Kaas, S. Kennedy, K. Kristiansen, J. R. Kultima, P. Léonard, F. Levenez, O. Lund, B. Moumen, D. Le Paslier, N. Pons, O. Pedersen, E. Prifti, J. Qin, J. Raes, S. Sørensen, J. Tap, S. Tims, D. W. Ussery, T. Yamada, H. B. Nielsen, M. Almeida, A. S. Juncker, S. Rasmussen, J. Li, S. Sunagawa, D. R. Plichta, L. Gautier, A. G. Pedersen, E. Le Chatelier, E. Pelletier, I. Bonde, T. Nielsen, C. Manichanh, M. Arumugam, J.-M. Batto, M. B. Quintanilha dos Santos, N. Blom, N. Borruel, K. S. Burgdorf, F. Boumezbeur, F. Casellas, J. Doré, P. Dworzynski, F. Guarner, T. Hansen, F. Hildebrand, R. S. Kaas, S. Kennedy, K. Kristiansen, J. R. Kultima, P. Leonard, F. Levenez, O. Lund, B. Moumen, D. Le Paslier, N. Pons, O. Pedersen, E. Prifti, J. Qin, J. Raes, S. Sørensen, J. Tap, S. Tims, D. W. Ussery, T. Yamada, P. Renault, T. Sicheritz-Ponten, P. Bork, J. Wang, S. Brunak, S. D. Ehrlich, A. Jamet, A. Mérieux, A. Cultrone, A. Torrejon, B. Quinquis, C. Brechot, C. Delorme, C. M’Rini, W. M. de Vos, E. Maguin, E. Varela, E. Guedon, F. Gwen, F. Haimet, F. Artiguenave, G. Vandemeulebrouck, G. Denariaz, G. Khaci, H. Blottière, J. Knol, J. Weissenbach, J. E. T. van Hylckama Vlieg, J. Torben, J. Parkhill, K. Turner, M. van de Guchte, M. Antolin, M. Rescigno, M. Kleerebezem, M. Derrien, N. Galleron, N. Sanchez, N. Grarup, P. Veiga, R. Oozeer, R. Dervyn, S. Layec, T. Bruls, Y. Winogradski, Z. Erwin G P. Renault, T. Sicheritz-Ponten, P. Bork, J. Wang, S. Brunak, S. D. Ehrlich and H. I. T. C. Meta (2014). “Identification and assembly of genomes and genetic elements in complex metagenomic samples without using reference genomes.” Nature Biotechnology 32(8): 822–828.

Nurk, S., D. Meleshko, A. Korobeynikov and P. A. Pevzner (2017). “metaSPAdes: a new versatile metagenomic assembler.” Genome research 27(5): 824–834.

Overbeek, R., R. Olson, G. D. Pusch, G. J. Olsen, J. J. Davis, T. Disz, R. A. Edwards, S. Gerdes, B. Parrello, M. Shukla, V. Vonstein, A. R. Wattam, F. Xia and R. Stevens (2014). “The SEED and the Rapid Annotation of microbial genomes using Subsystems Technology (RAST).” Nucleic Acids Res 42(Database issue): D206–214.

Parks, D. H., M. Imelfort, C. T. Skennerton, P. Hugenholtz and G. W. Tyson (2015). “CheckM: assessing the quality of microbial genomes recovered from isolates, single cells, and metagenomes.” Genome Research 25(7): 1043–1055.

Parks, D. H., C. Rinke, M. Chuvochina, P.-A. Chaumeil, B. J. Woodcroft, P. N. Evans, P. Hugenholtz and G. W. Tyson (2017). “Recovery of nearly 8,000 metagenome-assembled genomes substantially expands the tree of life.” Nature Microbiology 2(11): 1533–1542.

Pasolli, E., F. Asnicar, S. Manara, M. Zolfo, N. Karcher, F. Armanini, F. Beghini, P. Manghi, A. Tett, P. Ghensi, M. C. Collado, B. L. Rice, C. DuLong, X. C. Morgan, C. D. Golden, C. Quince, C. Huttenhower and N. Segata (2019). “Extensive Unexplored Human Microbiome Diversity Revealed by Over 150,000 Genomes from Metagenomes Spanning Age, Geography, and Lifestyle.” Cell 176(3): 649-662.e620.

Pride, D. T., R. J. Meinersmann, T. M. Wassenaar and M. J. Blaser (2003). “Evolutionary implications of microbial genome tetranucleotide frequency biases.” Genome Res 13(2): 145–158.

Quail, M. A., I. Kozarewa, F. Smith, A. Scally, P. J. Stephens, R. Durbin, H. Swerdlow and D. J. Turner (2008). “A large genome center’s improvements to the Illumina sequencing system.” Nature methods 5(12): 1005–1010.

Quince, C., A. W. Walker, J. T. Simpson, N. J. Loman and N. Segata (2017). “Shotgun metagenomics, from sampling to analysis.” Nature biotechnology 35(9): 833–844.

Raghoebarsing, A. A., A. Pol, K. T. Van de Pas-Schoonen, A. J. Smolders, K. F. Ettwig, W. I. C. Rijpstra, S. Schouten, J.S.S. Damsté, H. J. O. den Camp and M. S. Jetten (2006). “A microbial consortium couples anaerobic methane oxidation to denitrification.” Nature 440(7086): 918–921.

Ranjard, L., F. Poly and S. Nazaret (2000). “Monitoring complex bacterial communities using culture-independent molecular techniques: application to soil environment.” Res Microbiol 151(3): 167–177.

Rausch, P., M. Rühlemann, B. M. Hermes, S. Doms, T. Dagan, K. Dierking, H. Domin, S. Fraune, J. von Frieling, U. Hentschel, F.-A. Heinsen, M. Höppner, M. T. Jahn, C. Jaspers, K. A.B. Kissoyan, D. Langfeldt, A. Rehman, T. B. H. Reusch, T. Roeder, R. A. Schmitz, H. Schulenburg, R. Soluch, F. Sommer, E. Stukenbrock, N. Weiland-Bräuer, P. Rosenstiel, A. Franke, T. Bosch and J. F. Baines (2019). “Comparative analysis of amplicon and metagenomic sequencing methods reveals key features in the evolution of animal metaorganisms.” Microbiome 7(1): 133.

Rhoads, A. and K. F. Au (2015). “PacBio sequencing and its applications.” Genomics, proteomics & bioinformatics 13(5): 278–289.

Ribeca, P. and G. Valiente (2011). “Computational challenges of sequence classification in microbiomic data.” Briefings in bioinformatics 12(6): 614–625.

Sandberg, R., G. Winberg, C. I. Bränden, A. Kaske, I. Ernberg and J. Cöster (2001). “Capturing whole-genome characteristics in short sequences using a naïve Bayesian classifier.” Genome Res 11(8): 1404–1409.

Schloss, P. D. and J. Handelsman (2004). “Status of the microbial census.” Microbiology and molecular biology reviews 68(4): 686–691.

Sedlar, K., K. Kupkova and I. Provaznik (2017). “Bioinformatics strategies for taxonomy independent binning and visualization of sequences in shotgun metagenomics.” Computational and Structural Biotechnology Journal 15: 48–55.

Segata, N., D. Börnigen, X. C. Morgan and C. Huttenhower (2013). “PhyloPhlAn is a new method for improved phylogenetic and taxonomic placement of microbes.” Nature communications 4: 2304.

Sieber, C. M., A. J. Probst, A. Sharrar, B. C. Thomas, M. Hess, S. G. Tringe and J. F. Banfield (2018). “Recovery of genomes from metagenomes via a dereplication, aggregation and scoring strategy.” Nature microbiology 3(7): 836.

Strambio-De-Castillia, C., M. Niepel and M. P. Rout (2010). “The nuclear pore complex: bridging nuclear transport and gene regulation.” Nature Reviews Molecular Cell Biology 11(7): 490–501.

Sun, J., X.-P. Liao, A. W. D’Souza, M. Boolchandani, S.-H. Li, K. Cheng, J. L. Martínez, L. Li, Y.-J. Feng and L.-X. Fang (2020). “Environmental remodeling of human gut microbiota and antibiotic resistome in livestock farms.” Nature communications 11(1): 1–11.

Teeling, H., J. Waldmann, T. Lombardot, M. Bauer and F.O. Glöckner (2004). “TETRA: a web-service and a stand-alone program for the analysis and comparison of tetranucleotide usage patterns in DNA sequences.” BMC bioinformatics 5(1): 163.

Tettelin, H., V. Masignani, M. J. Cieslewicz, C. Donati, D. Medini, N. L. Ward, S. V. Angiuoli, J. Crabtree, A. L. Jones and A. S. Durkin (2005). “Genome analysis of multiple pathogenic isolates of Streptococcus agalactiae: implications for the microbial “pan-genome”.” Proceedings of the National Academy of Sciences 102(39): 13950–13955.

Tishkoff, S. A. and B. C. Verrelli (2003). “Patterns of human genetic diversity: implications for human evolutionary history and disease.” Annual review of genomics and human genetics 4(1): 293–340.

Tremblay, J., K. Singh, A. Fern, E. S. Kirton, S. He, T. Woyke, J. Lee, F. Chen, J. L. Dangl and S. G. Tringe (2015). “Primer and platform effects on 16S rRNA tag sequencing.” Frontiers in microbiology 6: 771.

Tringe, S. G., C. Von Mering, A. Kobayashi, A. A. Salamov, K. Chen, H. W. Chang, M. Podar, J. M. Short, E. J. Mathur and J. C. Detter (2005). “Comparative metagenomics of microbial communities.” Science 308(5721): 554–557.

Tyson, G. W., J. Chapman, P. Hugenholtz, E. E. Allen, R. J. Ram, P. M. Richardson, V. V. Solovyev, E. M. Rubin, D. S. Rokhsar and J. F. Banfield (2004). “Community structure and metabolism through reconstruction of microbial genomes from the environment.” Nature 428(6978): 37–43.

Uritskiy, G. V., J. DiRuggiero and J. Taylor (2018). “MetaWRAP—a flexible pipeline for genome-resolved metagenomic data analysis.” Microbiome 6(1): 158.

van der Walt, A. J., M. W. van Goethem, J.-B. Ramond, T. P. Makhalanyane, O. Reva and D. A. Cowan (2017). “Assembling metagenomes, one community at a time.” BMC Genomics 18(1): 521.

Vanwonterghem, I., P. D. Jensen, K. Rabaey and G. W. Tyson (2016). “Genome_Jcentric resolution of microbial diversity, metabolism and interactions in anaerobic digestion.” Environmental microbiology 18(9): 3144–3158.

Venter, J. C., M. D. Adams, E. W. Myers, P. W. Li, R. J. Mural, G. G. Sutton, H. O. Smith, M. Yandell, C. A. Evans and R. A. Holt (2001). “The sequence of the human genome.” science 291(5507): 1304–1351.

Venter, J. C., K. Remington, J. F. Heidelberg, A. L. Halpern, D. Rusch, J. A. Eisen, D. Wu, I. Paulsen, K. E. Nelson and W. Nelson (2004). “Environmental genome shotgun sequencing of the Sargasso Sea.” science 304(5667): 66–74.

Villegas, V. E. and P. G. Zaphiropoulos (2015). “Neighboring Gene Regulation by Antisense Long Non-Coding RNAs.” International Journal of Molecular Sciences 16(2): 3251–3266.

von Meijenfeldt, F. A. B., K. Arkhipova, D. D. Cambuy, F. H. Coutinho and B. E. Dutilh (2019). “Robust taxonomic classification of uncharted microbial sequences and bins with CAT and BAT.” bioRxiv: 530188.

Walsh, A. M., F. Crispie, O. O’Sullivan, L. Finnegan, M. J. Claesson and P. D. Cotter (2018). “Species classifier choice is a key consideration when analysing low-complexity food microbiome data.” Microbiome 6(1): 1–15.

Wang, Y., H. C. Leung, S.-M. Yiu and F. Y. Chin (2012). “MetaCluster 5.0: a two-round binning approach for metagenomic data for low-abundance species in a noisy sample.” Bioinformatics 28(18): i356–i362.

Whitaker, R. J. and J. F. Banfield (2006). “Population genomics in natural microbial communities.” Trends in Ecology & Evolution 21(9): 508–516.

Woloszynek, S., Z. Zhao, G. Ditzler, J. R. Price, E. R. Reichenberger, Y. Lan, J. Chen, J. Earl, S. K. Langroodi and G. Ehrlich (2018). Analysis Methods for Shotgun Metagenomics. Theoretical and Applied Aspects of Systems Biology, Springer: 71–112.

Wong, M. T., D. Zhang, J. Li, R. K. H. Hui, H. M. Tun, M. S. Brar, T.-J. Park, Y. Chen and F. C. Leung (2013). “Towards a metagenomic understanding on enhanced biomethane production from waste activated sludge after pH 10 pretreatment.” Biotechnology for biofuels 6(1): 1–14.

Woyke, T., H. Teeling, N. N. Ivanova, M. Huntemann, M. Richter, F. O. Gloeckner, D. Boffelli, I. J. Anderson, K. W. Barry and H. J. Shapiro (2006). “Symbiosis insights through metagenomic analysis of a microbial consortium.” Nature 443(7114): 950–955.

Wrighton, K. C., C. J. Castelle, M. J. Wilkins, L. A. Hug, I. Sharon, B. C. Thomas, K. M. Handley, S. W. Mullin, C. D. Nicora and A. Singh (2014). “Metabolic interdependencies between phylogenetically novel fermenters and respiratory organisms in an unconfined aquifer.” The ISME journal 8(7): 1452–1463.

Wu, Y.-W., B. A. Simmons and S. W. Singer (2015). “MaxBin 2.0: an automated binning algorithm to recover genomes from multiple metagenomic datasets.” Bioinformatics 32(4): 605–607.

Wu, Y.-W. and Y. Ye (2011). “A novel abundance-based algorithm for binning metagenomic sequences using l-tuples.” Journal of Computational Biology 18(3): 523–534.

Wyrick, J. J. and R. A. Young (2002). “Deciphering gene expression regulatory networks.” Current Opinion in Genetics & Development 12(2): 130–136.

Yang, B., Y. Peng, H. C. Leung, S.-M. Yiu, J. Qin, R. Li and F. Y. Chin (2010). MetaCluster: unsupervised binning of environmental genomic fragments and taxonomic annotation. Proceedings of the first ACM international conference on bioinformatics and computational biology.

Young, A. L., H. O. Abaan, D. Zerbino, J. C. Mullikin, E. Birney and E. H. Margulies (2010). “A new strategy for genome assembly using short sequence reads and reduced representation libraries.” Genome research 20(2): 249–256.

